# Nutritional stress alone can control *in vitro* tumor formation and its invasive nature

**DOI:** 10.1101/2020.02.17.952234

**Authors:** Sukanya Gayan, Abhishek Teli, Anish Nair, Tuli Dey

## Abstract

The metastatic nature is an inherent property of the tumor. However, the effect of the environmental stress conditions on tumor microcosm in the context of metastasis needs to be analyzed. This work is proposed to analyze the tumor behavior under multiple metabolic stress conditions, such as deprivation of glucose, protein, and oxygen. The spheroid proliferation rate is observed to be influenced profoundly by the stress level where minimal stress produces compact spheroid and severe stress makes unstable aggregate like structures. It is observed that the non-invasive cancer cells cannot form spheroids under extreme stress. Stress conditions influence the mRNA levels of hypoxic, angiogenic and ECM deformation specific gene clusters. Spheroid reversal assay reveals the quiescent nature of the stressed spheroids under continuous stress conditions. However, after the rescue, the stressed spheroids were found to opt for different migration modalities. Extremely stressed non-invasive spheroids display atypical sprout-like growth within the invasion matrix and severely stressed spheroids can control the migration pattern of mesenchymal stem cells. Thus, it is concluded that multiple nutritional stress influences the spheroid formation and physiology along with the conversion of a non-invasive spheroid into quasi-invasive one.

## Introduction

Cancer is one of the most fatal diseases due to its metastatic behavior. Tumors (>2-5 cm diameter) with a necrotic core, and a proliferating periphery initiate the shedding of metastatic cells and instigate the growth of occult tumors in other tissues ^1^. Migration of tumor cells towards the greener pasture of future metastatic niches is a complex process. Therefore, understanding of metastasis at a molecular level has gathered much interest, because of its potential to provide much deeper insight into the tumor’s social intelligence and probable cure ^2^. Many factors such as tumor microenvironment, extra cellular matrix and the environmental stress are presumed to influence metastasis ^3, 4^. Additionally inter-cellular conditions like deprivation of nutrition and oxygen are assumed to play influential role in angiogenesis and aggression of the tumor ^5, 6^. Rapidly growing tumors utilize the available glucose inefficiently and initiate acidosis ^7, 8^. Experimental deprivation of glucose is found to induce cell death *in vitro* while in xenograft studies it has been found to activate AMPK and ATF4 pathways to initiate angiogenesis ^9, 10^. Similarly protein or amino acid (such as arginine) starvation induces apoptosis which can be bypassed through the GCN2/ATF4 Pathway ^11, 12^. Hypoxic stress in cell culture condition induces apoptotic cell death and plays an important role in autophagy ^13-15^. However in tumors hypoxic stress has also been highlighted as one of the most important factor to initiate angiogenesis and metastasis ^16, 17^. In fact, this multifaceted relationship between angiogenesis and metastasis seems to be synergistic in nature, as anti-angiogenic therapies have been found to reduce both the incidence and severity of metastasis ^18, 19^. Reports of metastasis stimulated by the anti-angiogenic therapy further demonstrate the complicated nature of this relationship between angiogenesis and metastasis ^20, 21^.

In stark contrast to the underlying complexities between these phenomena, currently available *in vitro* model systems seem far too simplistic. Most of the models consider single stress condition while studying, whereas in a real dynamic scenario the tumor is never exposed to any individual stress, but actually to multiple stresses that have a collective effect. In spite of this critical limitation, to date only one study has attempted to understand the multi-stress conditions ^22^. The encouraging effect of nutrient stress on tumor metastasis or epithelial to mesenchymal transition (EMT) has been studied in the past ^23, 24^. Additionally, till date, the direct effect of metabolic stress has been studied mainly in two dimensional (2D) platforms, which can’t depict the complexity of 3D tumors. It has been assumed that multicellular tumors can sense the environmental stress and modulate its own transcriptional and translational machinery to either evade or tolerate the stress which cannot be mimicked by the cellular monolayer ^25, 26^. Though xenograft model is successfully used in this scenario, existing technical and analytical challenges make it a difficult and costly affair. Additionally, control of experimental variables and the ability to envision the concept within the *in vivo* model system seems to be very problematic ^27^. In contrast, the *in vitro* models such as multi-cellular spheroids can provide a middle ground, with more control over the many variable parameters for study.

The current study is aimed to be the first among such studies to fill the lacunae by identifying the effect of multiple stress conditions (such as individual or combined starvation of glucose, protein, and oxygen) on a multicellular non-invasive tumor model.

## Results

### 1. Creation of different level of nutrient stress

Four different stress levels such as minimal, moderate, severe, and extreme are created by combining different stress conditions (SI-Fig 1). Low stress conditions are created by depriving only nutrition molecules (such as glucose, serum and oxygen) individually, while moderate-severe stress is created by combining the deprivation of both nutrition and oxygen (nomenclature is mentioned in method and materials). By changing the exposure to the stress condition, further subtypes of stress have been created, such as acute and chronic. In the case of acute stress, spheroids are fabricated and exposed to stress thereafter. Under the chronic stress conditions, already stressed cells are used to fabricate spheroids. As observed, low to moderate stress (mostly acute) exhibit no deleterious effect on spheroid formation. However, severe to extreme stress conditions (mostly chronic) are found to control the spheroid formation *in vitro* and its morphology negatively.

### 2. Different stress conditions influence the spheroid growth and morphology differentially

From the phase contrast images, it is evident that individual nutritional and hypoxic stresses are not deterrent to spheroid formation even under acute and chronic conditions (Fig 1A, 3A). From the aspect ratio analysis, the acute nutrional stress condition especially L-0 found to shrink the spheroids significantly but L-5 increase the size (Fig 1B). Hypoxia alone exhibits no detrimental effect on spheroid morphology and growth. Chronic individual stress conditions show no visible change in spheroid morphology, but negatively influence the spheroid growth over time. As observed L-5 and L-0 conditions negatively affect the spheroid size (Fig 3A-B). Hypoxia in chronic condition shows no negative effect on spheroid formation and growth. Spheroids fabricated under individual acute and chronic conditions (nutritional and hypoxic) resemble the control morphologically with a well defined periphery and compact nature. On the other hand spheroid morphology is severely affected under acute combined stress condition (Fig 5A-B). Interestingly, the acute combined stress conditions such as H-0-Hy and L-0-Hy change the spheroid morphology and they appear as spheroids of invasive breast cancer cell line (MDA MB 231) exhibiting loosely assembled structures with irregular boundary. Spheroid size has increased significantly under H-0-Hy (Fig 5B).

**Figure 1.**
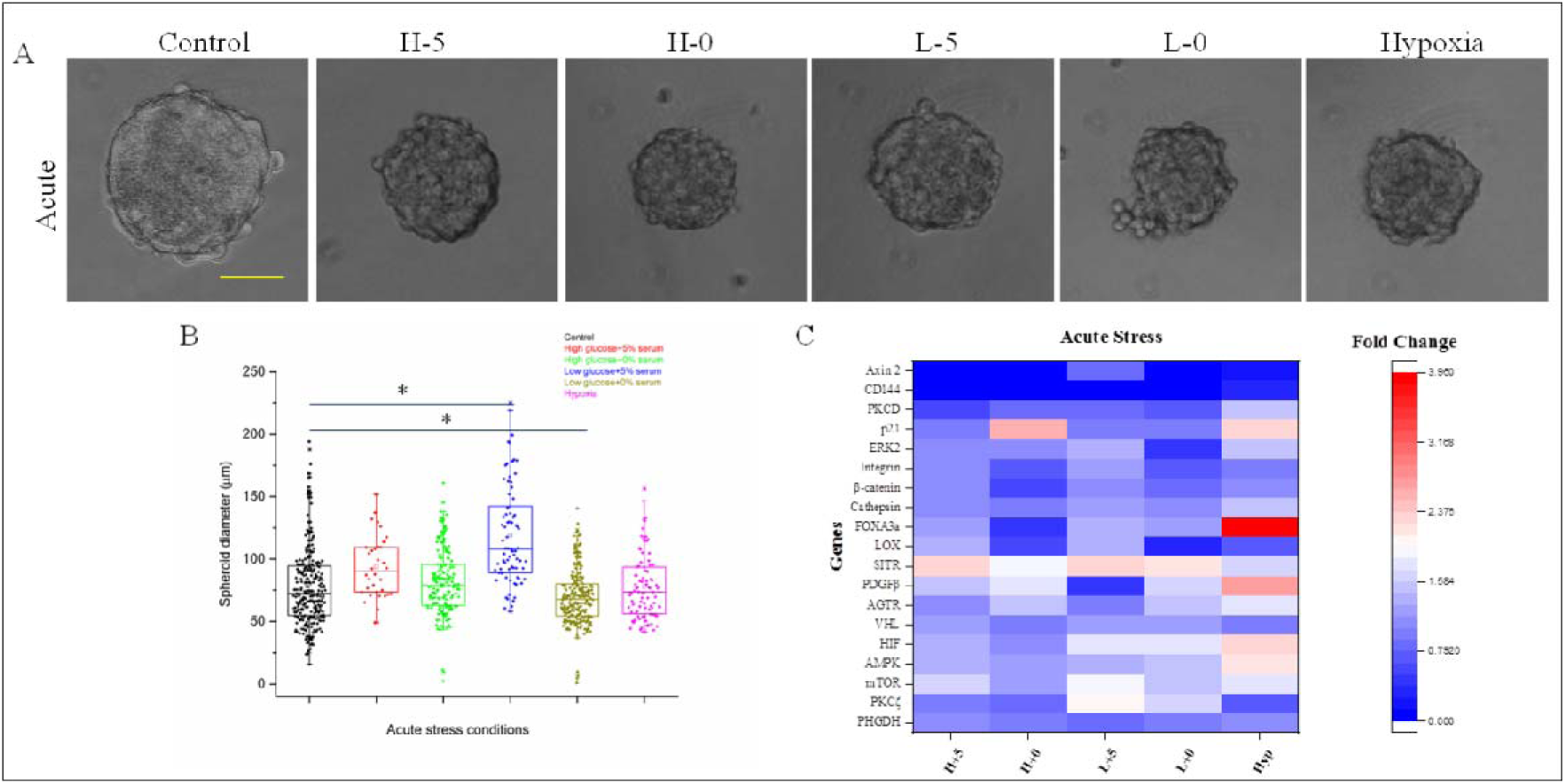
Effect of acute nutritional and hypoxic stress on spheroid formation and growth. A- Phase contrast image of control and spheroids under stress conditions. B- Spheroid size analysis under stress conditions. C- Gene expression profile of spheroid under stress conditions. Normalized fold change values were plotted using a heat map module. Scale bar is 100 µm.

**Figure 2.**
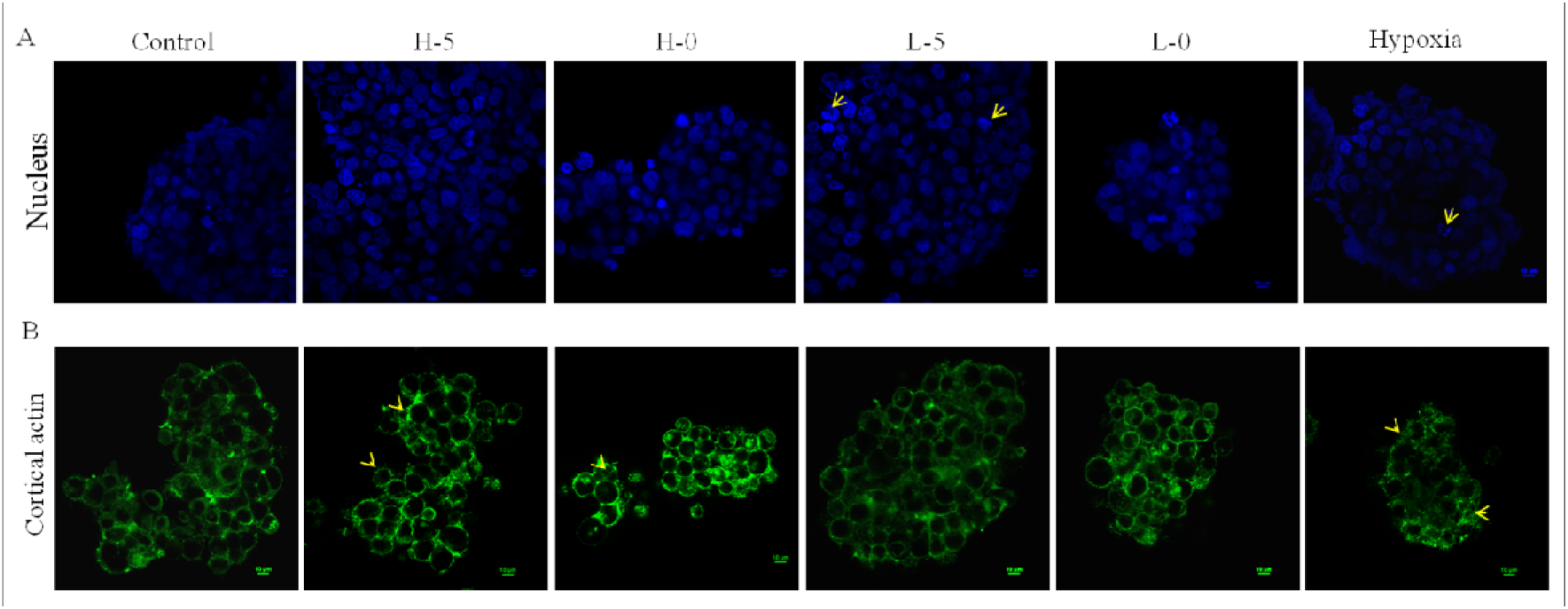
Effect of acute nutritional stress on nucleus morphology and cortical actin distribution. A- Morphological analysis of the nucleus within the control and stressed spheroids. Yellow arrows indicate apoptotic nuclei. B- Cytoneme distribution in stressed and control spheroids (green is used as pseudocolor). Spheroids were fixed, permeabilized and stained with Rodamine conjugated phalloidin followed by imaging in CLSM (60x objective). Scale bar was 10 µm.

**Figure 3.**
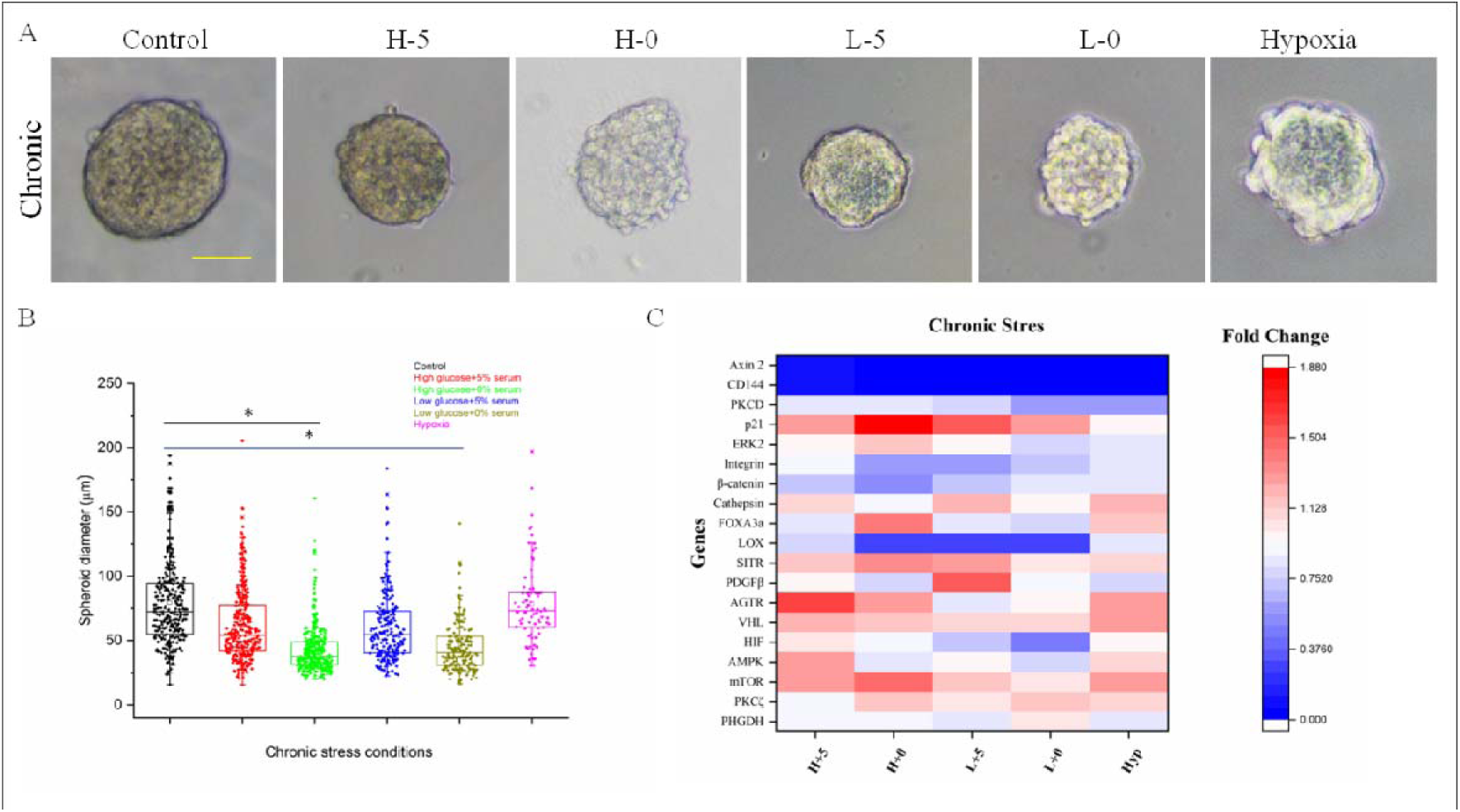
Effect of chronic nutritional and hypoxic stress on spheroid formation and growth. A- Phase contrast image of control and spheroid under stress conditions. B- Spheroid size analysis under stress conditions. C- Gene expression profile of spheroid under stress conditions. Normalized fold change values were plotted using a heat map module (origin lab). Scale bar is 100 µm

**Figure 4.**
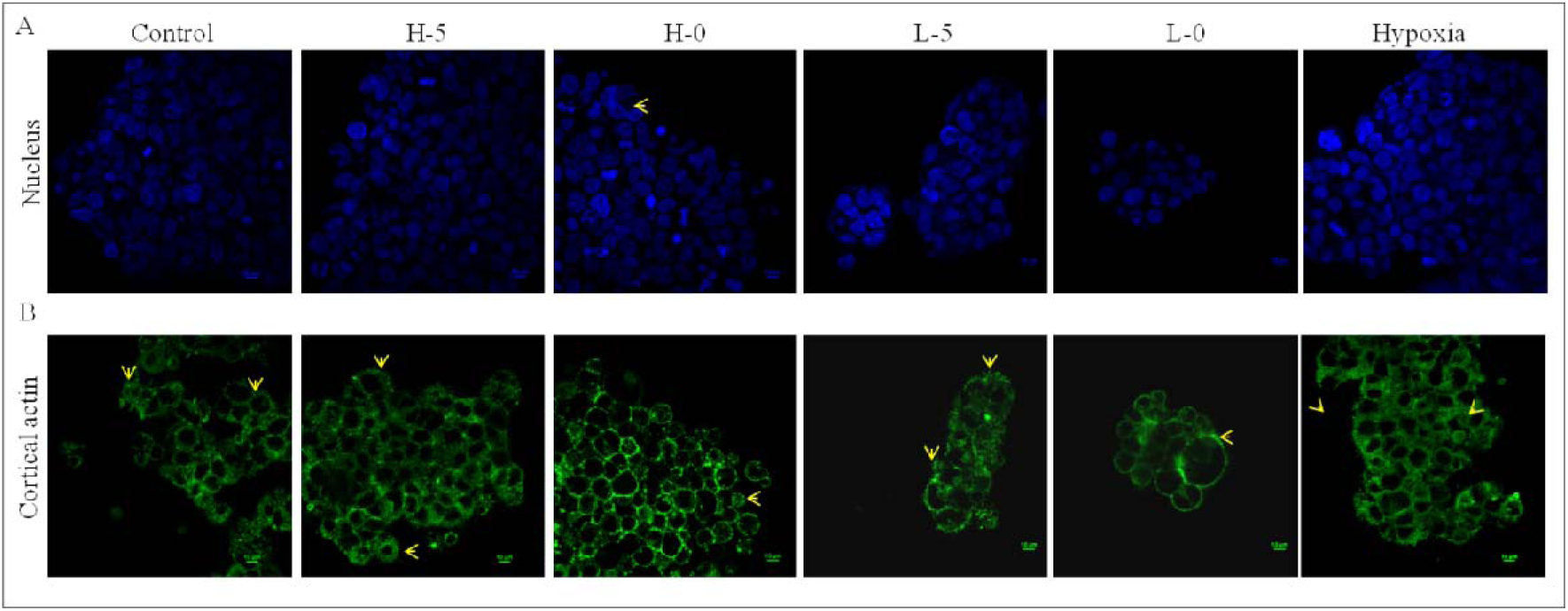
Effect of chronic nutritional and hypoxic stress nucleus morphology and cortical actin distribution. **A-** Morphological analysis of the nucleus within the control and stressed spheroids. Yellow arrows indicate apoptotic nuclei. B- Cytoneme distribution in stressed and control spheroids (green is used as pseudocolor). Spheroids were fixed, permeabilized and stained with Rodamine conjugated phalloidin followed by imaging in CLSM (60x objective). Scale bar was 10 µm.

**Figure 5.**
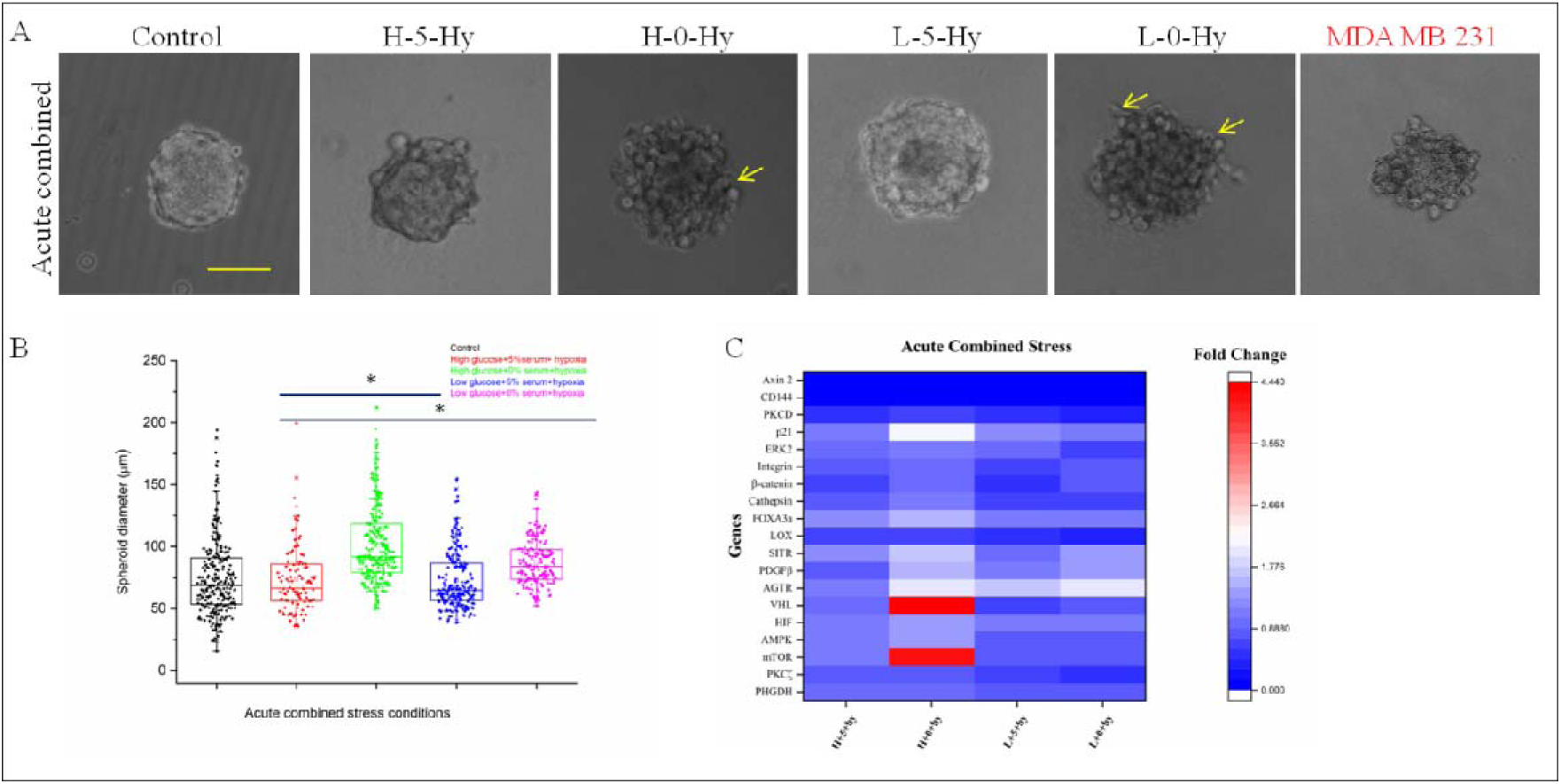
Effect of the acute combined stress (nutritional and hypoxic) on spheroid formation and growth. A- Phase contrast image of control and spheroid under stress conditions. Invasive spheroid of MDA MB 231 is marked in red. B- Spheroid size analysis under stress conditions. C- Gene expression profile of spheroids under stress conditions. Normalized fold change values were plotted using a heat map module. Scale bar was 100 µm.

The chronic combined stress inhibits the spheroid formation completely (Fig 6A). However, when Cultrex™ spheroid formation matrix is used, spheroids are observed to form under chronic combined stress probably due to the presence of ECM based factors. The morphology of fabricated spheroids highly resembled the invasive spheroid (MDA MB 231) (Fig 6A).

**Figure 6.**
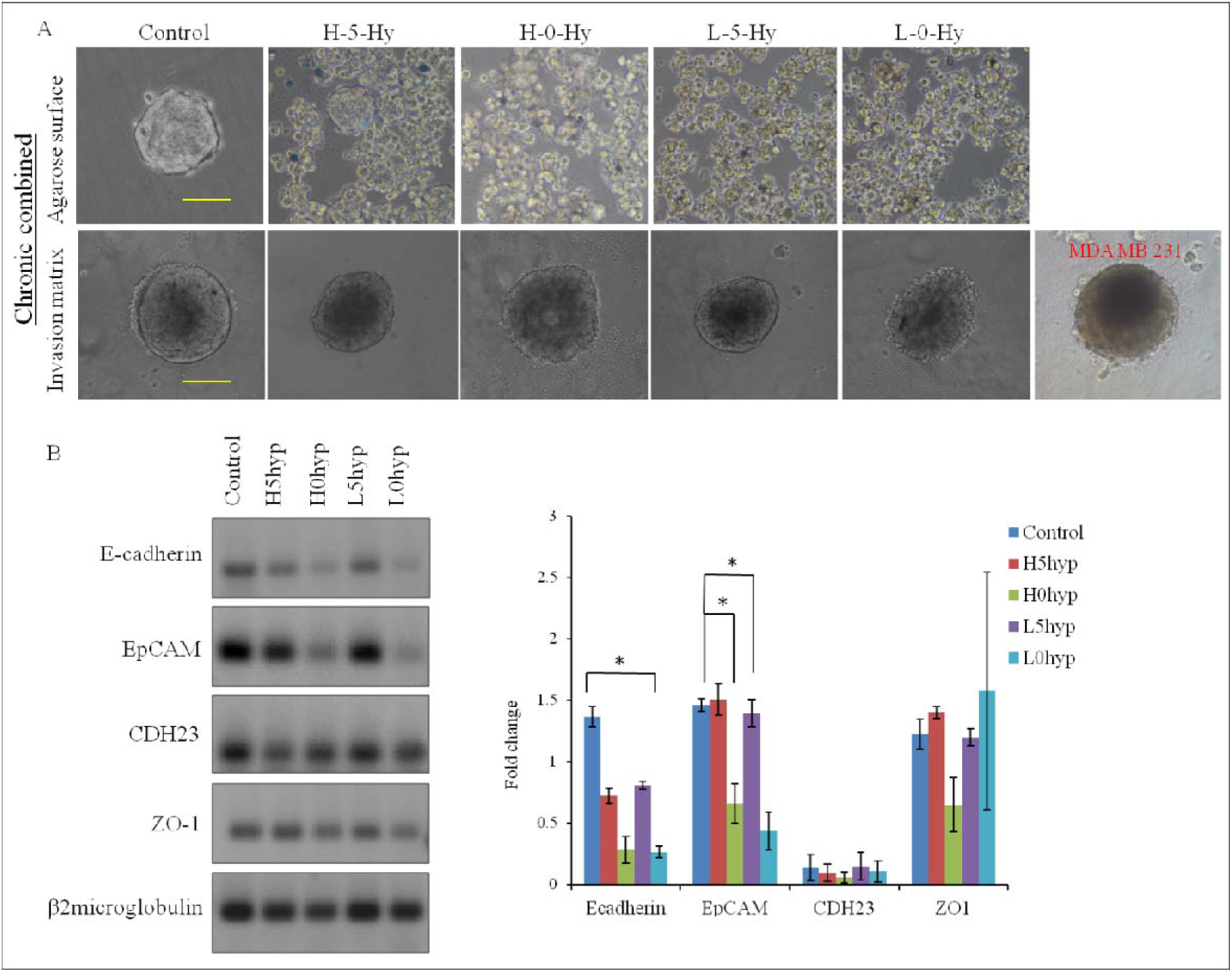
Effect of the chronic combined stress on spheroid formation. A- Phase contrast image of cells not aggregated under chronic combined stress conditions. B- Control and formed spheroid under similar stress conditions fabricated in the Cultrex™ invasion matrix. Invasive spheroid of MDA MB 231 fabricated under the same condition was marked in red. C- Gel electrophoretic data and densitometric analysis of cell adhesion specific mRNA expression followed by semi-quantitative RT-PCR. Scale bar was 100 µm.

Cell adhesion molecules (CAMs) are generally down-regulated in invasive (MDA MB 231) cells and they negatively influence the spheroid formation. Expression of some representative CAMs (E cadherin, EP-Cam, Zho 1 and Cad23) in chronic combined stressed spheroids are analysed by semi-quantitative RT-PCR. Our data suggests that chronic combined stress leads to the down-regulation of E-cad and Ep-CAM without any significant change in the levels of Zho1 and Cad23 (Fig 6B-C).

### 3. Nutritional stresses influenced the expression of genes involved in glycolysis, amino acid synthesis, angiogenesis, cancer stemness, hypoxia and matrix deformation pathways

In our study, individual and combined stresses in acute and chronic conditions are found to influence the mRNA level of genes involved in glycolysis, amino acid synthesis, angiogenesis, cancer stemness, hypoxia and matrix deformation (SI-Table 1). The acute individual stress up-regulates the mRNAs related to angiogenesis (SITR), protein starvation (P21, ERK 1/2) and matrix deformation (LOX, FOXA3a) pathway genes (Fig 1C), whereas in acute combined stress (H-0-Hy), only angiogenesis (VHL) and glycolysis shift (mTOR) genes get up-regulated significantly (Fig 2C). Other mRNAs are mostly down-regulated or show no significant change between the two conditions. In chronic individual condition, matrix deformation (FoxA3a, cathepsin), protein starvation (PKCD, P21) and angiogenic (SITR), hypoxic (VHL, HIF, AGTR) glycolysis shift (AMPK, mTOR) clusters exhibit significant increment compared to control (Fig 3C). Conversely, stemness related (Axin 2, CD144) and cell-ECM adhesion genes (integrin, β-catenin) showed significant down-regulation along with the CAM protein as mentioned earlier. Amongst all the stress levels, L-0 condition exhibit maximum transcriptional dormancy compared to others. Hypoxia, as an individual stress exhibits significant effect under chronic condition compared to the acute (Fig 1C-2C). To understand the global effect of this differential expression profile of mRNAs, the analysis mode of Reactome Knowledgebase (https://reactome.org) is used. Over-represented pathways have been highlighted in the genome-wide overview of the results of pathway analysis (SI Fig 2). Statistically significant pathways involving the candidate genes are enlisted in SI-Table 2.

### 4. Nutrional stress and its effect on the nucleus

Under both acute and chronic individual stress conditions, no significant difference in nucleus morphology is observed (Fig 2A, 4A). However the nuclear surface area of top 1-2 cell layers is observed to increase in acute hypoxia condition and decrease significantly in acute H-0 and L-0 conditions (SI Fig 3). Effect of chronic stress on nucleus size seems more prominent as H-0, L-5 and L-0 show significant decrease in size compared to hypoxic one. Interestingly, no such variation is observed in the middle layer nucleus of spheroids (data not shown). Additionally minimal apoptotic bodies are observed in acute stress conditions (such as L-5, L-0 and Hypoxia) (Fig 4A). Serum deprivation is observed to induce the ‘cell-in cell’ structure specifically in acute L-5 and chronic H-0 condition (SI-Fig 4).

### 5. Nutrional stress control cortical actin distribution and cytoneme formation

Distribution of actin cytoskeleton is observed to be cortical in both acute and chronic stress conditions. However, the thickness of cortical actin ring is seemed to be decreased under acute and chronic stress conditions (SI Fig 5). Additionally in severe stress conditions, the peripheral cells showed increased amount of actin-rich membrane protrusions or cytoneme as compared to the control spheroids.

### 6. Continuous stress induces dormancy

Migratory behavior of the spheroids under different conditions is described in tabulated form (SI-Table 3). Under favourable conditions (control) the MCF7 spheroid pursues epithelial (collective) migration. In the case of continuous stress most of the spheroids exhibits no migration at all, while display epithelial migration (SI-Fig 6).

### 7. Rescued spheroids induced single cell migration and sprouting in non-invasive spheroids

When rescued, an interesting profile is seemed to emerge where the rescued spheroids display different types of migration. Moderately stressed spheroids (acute H-5, H-0, L-5, Hypoxia and chronic H-5, L-5, Hypoxia) show epithelial migration as control spheroids. Severely stressed spheroids shows only single cell migration (acute L-0) and mixed mode of migration (H-5-Hy, H-0-Hy, and L-5-Hy) while extremely stressed spheroids (chronic H-0, L-0, and acute combined L-0-Hy) remain complete dormant (Fig 7A-8A). The single cell migration inducing conditions (acute L-0, H-5-Hy, H-0-Hy, and L-5-Hy) are further investigated to identify the mode of migration. They exhibit low expression of MMPs (SI-Fig 7A) and can’t be inhibited by marimastat (MMP inhibitor) but by ML7 (myosin light chain kinase inhibitor) (SI-Fig 7B). Chemo-tactic migration of spheroids from chronic combined stress is carried in Cultrex™ invasion matrix (Fig 8B). The continuously stressed spheroids remain dormant (data not shown), while the rescued spheroids exhibit the initiation of sprout like projections, which are not visible in control or in any moderately stressed spheroids (H-5-Hy).

**Figure 7.**
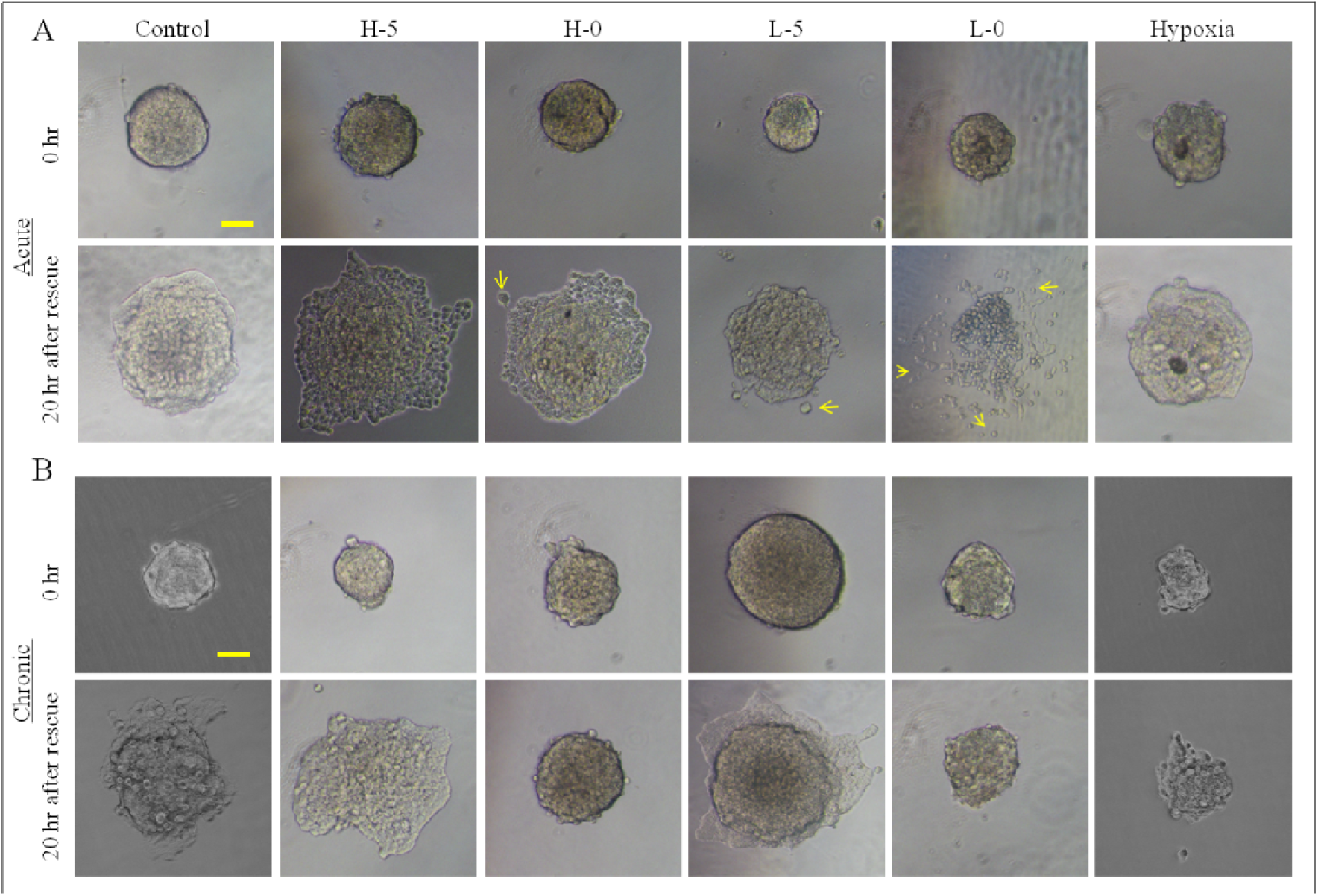
Pseudo 3D migration profiling of stressed (A-acute individual, B-chronic individual) spheroids after the rescue. Stressed spheroids were treated with fresh cell culture media and incubated in standardized cell culture conditions for 20-24 hr. Imaging was done in different time points using phase-contrast microscope. Yellow arrows indicate a single cell. Scale bar was 100 µm.

**Figure 8.**
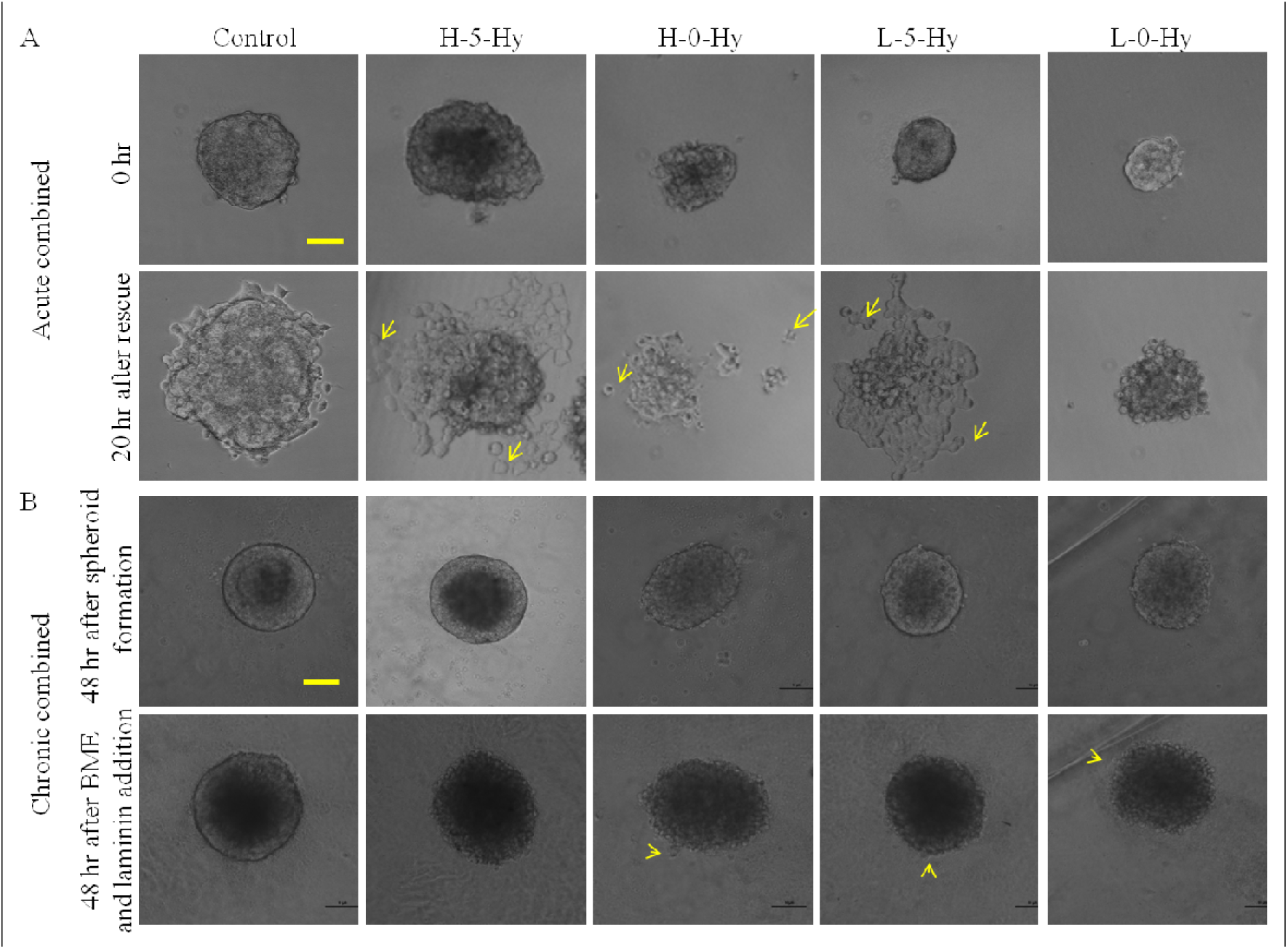
Pseudo 3D migration profiling of stressed (A-acute combined, B-chronic combined) spheroids after the rescue. Stressed spheroids were treated with fresh cell culture media and incubated in standardized cell culture conditions for 20-24 hr. Imaging was done in different time points using phase-contrast microscope. Yellow arrows indicate a single cell or sprouts. Scale bar was 100 µm.

### 8. Severely stressed spheroids influence stem cell migration after rescue

After rescue, severely stressed spheroids (H-5-Hy, H-0-Hy, L-5-Hy and L-0-Hy) (green) are found to influence the stem cell (red) migration. It is observed that within 24 hr, the spheroid induces a directional migration of stem cells toward the spheroid (Fig 9). On the control sample (without spheroid) the stem cells remain scattered throughout the gel surface, and in the case of control spheroid the stem cells penetrate the gel and create a ring like structure surrounding the spheroid. With increasing stress (H-5-Hy<L-5-Hy<H-0-Hy<L-0-Hy) the circles from top become more concentric. Increased penetration of the gel by the stem cells is also observed in higher stress conditions such as H-0-Hy, L5-Hy, and L-0-Hy. Recruitment of stem cells within the stressed spheroid structure is also significantly higher compared to the control spheroid.

**Figure 9.**
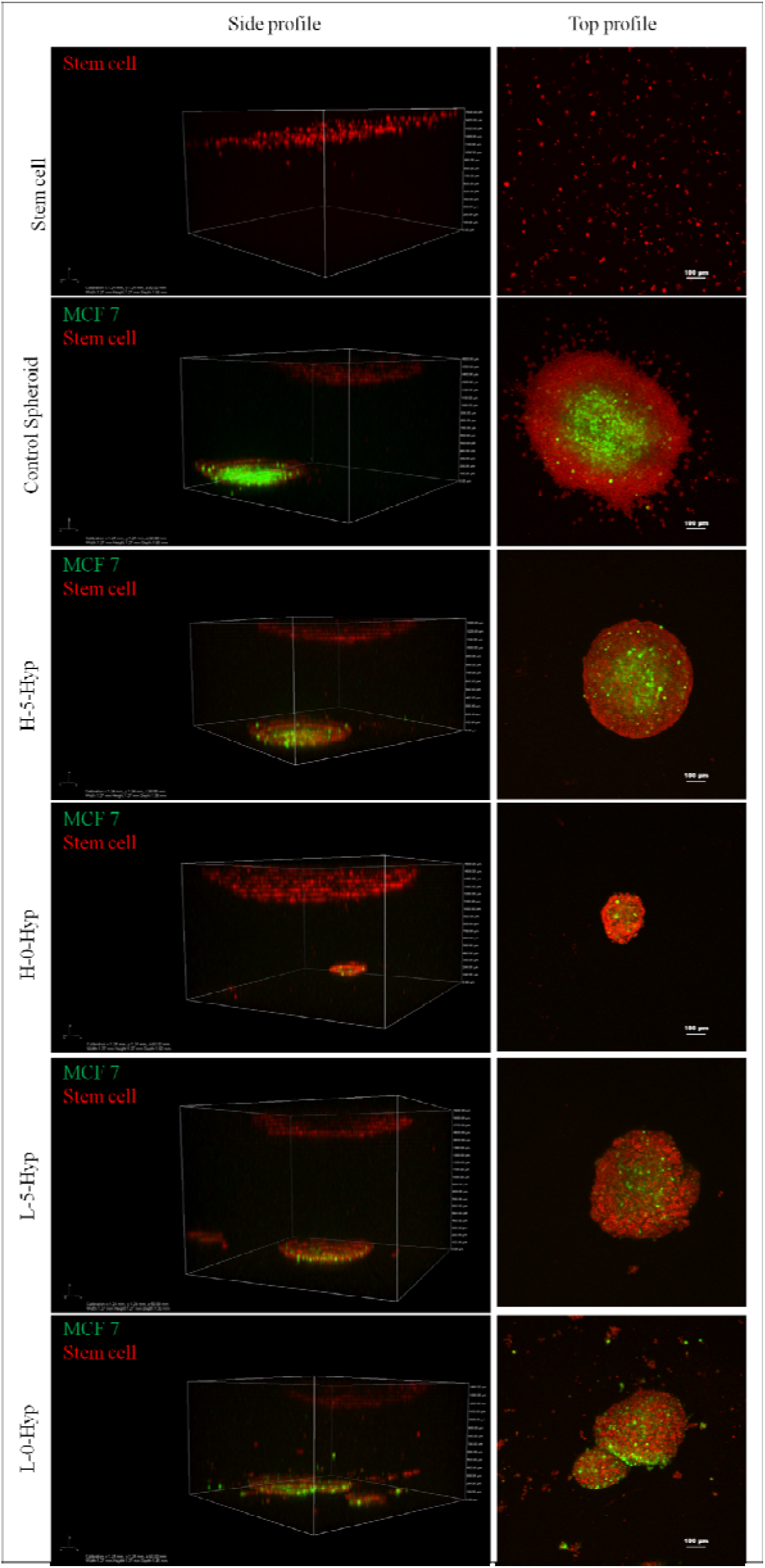
Directional migration of stem cells under the influence of stressed spheroids. Control and stressed spheroids were encapsulated in collagen gel followed by the seeding of Human mesenchymal stem cells (pre-stained with Cytotracker Red™) on top of it. Imaging was done after 72 hr with CLSM (10x objective). Side profile analysis was done with NIS Element software and presented as left panel. The distance between the top and bottom layers is 1000-1300 µm. The yellow arrows were indicating the spheroid at the bottom. The top view of the spheroid is presented as right panel.

## Discussion

During the preliminary growth phase, tumor cells show unprecedented proliferation and excessive appetite for nutrition. However, the inefficient utilization of available glucose through lactic acidosis can create glucose deprivation ^7, 8^. Increasing size of tumors can create oxygen deprivation within, due to the unavailability of blood capillaries which further increases the deficiency ^25^. Influence of such stresses on tumor behavior is very important, as it might manipulate the growth and progression of the tumor ^5, 16, 19^. A tumor, as a smart enterprise, has the potential to circumvent the nutritional stress ^6, 25, 28^ or let the microenvironment dictates its fate ^29^. The concept of movement ecology has initiated a new understanding of cancer as an invasive species, where it has intrinsic motivations to move or to stay. The decision of migration is mostly depends on the native ecosystem, where the deteriorating condition might put the organism’s survival at risk ^30^. The metabolic heterogeneity of the tumor microenvironment probably paves the way for deregulated metabolic properties, to survive hypoxia and glucose deprivation ^31^. We hypothesized that tumor behavior can be predicted, if we can understand the relationship between the metabolic stress and tumor’s response.

The current study is designed to understand the collective effect of metabolic stress on the tumor behavior through a multicellular spheroid based model. We have created different levels of stress for the spheroid. To create the ‘acute’ (minimal) stress, the spheroids are fabricated first and then exposed to stress conditions for 72 hr. Parallelly monolayer cancer cells are exposed to ‘chronic’ (severe) stress and used afterward to fabricate spheroids. We assumed that cancer cells will perceive the level of stress and respond accordingly. As expected the severely stressed cells observed to form the spheroids mimicking the invasive spheroids (MDA MB 231 spheroids) visually. But the mono-layer cells exposed to the combined stress (extreme), don’t form any spheroids. It can be assumed that after the multicellular structures are formed, deprivation of individual factors is not enough to resist the growth. However, chronic stress conditions (such as H-5, H-0, L-5 and L-0) indicate the dire effect of nutrition withdrawal during the formation of tumor. Hypoxia, as an individual stress has no visible impact on spheroid formation and growth either in acute or chronic mode, which exhibits the tolerance of multicellular structure towards oxygen deprivation. Failure of glucose and serum starvation to induce any deleterious effect, further strengthen the assumption that tumor physiology responds differently. Additionally we have observed that acute L-5 and acute H-0-Hy conditions exhibit a statistically significant increment in spheroid size, which can be explained by the auto-activating mutations of Ack1 of cancerous cells followed by the Tyr176 and Ser473/Thr308 phosphorylation of AKT under serum starvation. This AKT activation arc further maintain the cellular proliferation under stress conditions ^33, 34^. However complete removal of serum (L-0) restricts the proliferation. The observation that extreme stress condition (minimal nutrient) can hinder the spheroid formation completely is in line with the recent report of finding lowest BMI population with lowest risk of cancer ^35^.

Failure to form stable cellular aggregation (tumor) under extreme stress conditions can also explained by the reduced mRNA level of E-cad and EpCAM. Reduced level of homotypic cell adhesion molecules are in line with previous observation in the aggressive cell line (MDA MB 231) ^36, 37^. Visually the control and minimally stressed spheroids exhibit a well defined boundary and compactness which is altered in extreme stress conditions as predicted earlier ^29^. The irregular shape and loose assembly, highly resembles the invasive spheroids created *in vitro* (MDA MB 231) ^38^ and cancerous masses ^39, 40^. It can be assumed that the nutritional stress alone have the capacity to reprogram the original fate of the non-invasive spheroid into an invasive one. This might indicate the adaptable nature of the stressed cells according to the micro-environment.

Analysis of nucleus morphology reveals that peripheral cells exhibit an arising trend of flattened morphology compared to control. Swelling of the hypoxic nuclei without losing the integrity of nuclear envelope is in line with previous report ^41^. Nutritional and hypoxic stress has been found to initiate apoptosis in two dimensional culture conditions but very few apoptotic nuclei are observed here which can be explained by the anti-apoptotic nature of the spheroid structures ^43^. Serum starved spheroids exhibit multi-nucleation which might have been caused by the mitotic stress on cell cycle or entosis ^44, 45^. Cell-in-cell phenomena are observed in acute L-5 and chronic H-0, which might highlight the survival mechanism of ‘eating neighbours’ observed in the tumors under stress ^45^.

mRNA profile of glycolysis, amino acid synthesis, angiogenesis, stemness, matrix deformation gene cluster exhibit a common profile, where the acute stress (both individual and combined) down-regulate most of the genes, creating a transcriptomically dormant state. However, chronic individual stress tends to up regulate most of the genes from matrix deformation, protein starvation, angiogenic, hypoxic and glycolysis shift cluster thus exhibiting the urge to fight for survival. Stemness cluster shows no change in any case, which can be explained by the assumption that the experimental exposure here is not enough to activate the stem cell properties. Further analysis through ‘Reactome Knowledgebase’ shows over-representation of pathways related to signal transduction, Cell cycle, apoptosis, autophagy, transcription and extracellular matrix organization^46^.

Prominent cellular behavior such as migration, cell division, etc is controlled by the cytoskeletal machinery. Minimal-extreme metabolic stress exhibits no significant difference on actin distribution. However with increasing stress, the cortical actin thickness tends to decrease which might influence the cellular contractility and stability ^47^. Additionally the metabolic stress conditions were found to increase the membrane ruffling and formation of actin rich macropinocytosis. Abundance of macropinocytes or cytoneme like structure in stressed spheroids might control the entosis or nutrient-scavenging mechanisms to support the tumorogenesis within the nutrient-deprived environments ^48^.

Metabolic stress conditions are found to influence the mode of migration of multi-cellular spheroid which can also act as an indicator of altered behavior and emerging invasive nature. Diverse migration pattern is observed under stressed or rescued condition which highlights the newly observed plasticity of the non-invasive spheroids for the first time. Emergence of quiescent stage and minimal migration under continuous stress here is in line with the previous reports of tumor dormancy ^49^. The awakening of dormant cells through nutritional rescue is of importance here as rescued spheroids exhibit a spectrum of migration modalities ranging from collective (no-moderate stresses conditions), single cell (extreme individual stress) and mixed (extreme combined stress) migration. Non-invasive tumors generally use collective/epithelial migration. However when rescued from extreme stress conditions spheroids initiate single cell migration which might mimics the metastatic shedding. Diminished level of MMP secretion in those conditions (Acute L-0, H-5-Hy, H-0-Hy, and L-5-Hy) highlights the probability of amoeboid modality. Differential inhibitor assay shows the significance of MLC phosphorylation and highlight the role of MLCK induced amoeboid migration. It can be assumed that metabolically stressed spheroids prefer to choose amoeboid migration as an escape route ^17^. Chronic combined stressed spheroids exhibit single cell protrusions under the rescued condition within the invasion matrix, which is atypical of non-invasive cells such as MCF7, but observed in invasive spheroids ^36^.

Tumor growth and proliferation depends on the stromal cell population which might play an important role in healing stressed tumors ^50^. Direct co-culture invasion assay exhibits the tumor mediated directional migration and recruitment of stem cells as reported earlier. Among the stressed spheroids, only the severe ones (L-5-Hy, H-0-Hy and L-0-Hy) are found to control the migration of stem cells more profoundly compared to the control and H-5-Hy. It can be assumed that metabolically stressed tumors are more proficient in instructing the stem cells for metabolical support. ^51^.

The current study describes the cumulative effect of metabolic stress on an *in vitro* tumor model for the first time and can become instrumental in understanding the tumor behavior. Furthermore this report highlight the importance of nutrient availability and its effect on *in vitro* tumor formation which can be linked with the recent report on lowest BMI and lowest cancer risk ^35^. Metabolically challenged spheroids can serve as a platform for studying the synergistic effect of fasting and chemotherapy cycles, which proves to be more effective and therapeutically potent for wide range of tumors ^52^. It is observed here and supported by numerous other reports that individual stress conditions alone such as hypoxia is not enough to introduce *de novo* capability for organ colonization. But from the current study it can be concluded that multiple nutritional stress along with hypoxia can recondition the non-invasive spheroid into a quasi-invasive one, and it might have long term implications on the survival through the enhanced recruitment of stromal cells and growth of occult-tumors in distant tissues.

## Conclusion

Though genetic and epigenetic changes at global range play the most important role in triggering the metastatic cascade of cancer, local environmental stress is also crucial for the metastatic predisposition of cancer cells. Tumor cells are capable of redesigning their cellular pathways in order to resist or accept the micro-environmental changes such as hypoxia or nutritional stress, thereby exerting additional survival advantages. Additionally, the adverse environmental cues can influence the phenotype of the tumor and might aid in tumor cell plasticity towards its metastatic traits. Spheroids under combined stress (nutritional and hypoxic) exhibit such phenotypic plasticity in terms of migration and recruitment of stromal cells. In future these can be used as model to study the metabolical change that happen within the tumor cells and seems to influence the tumor-environment crosstalk.

## Methods and Material

### 1 Materials

DMEM with high glucose (4.5 gm/lt) and low glucose (1 gm/lt), fetal bovine serum, penicillin-streptomycin, Trypsin-EDTA solution and CoCl_2_ were procured from Himedia Laboratories, India. Trizol, Revertaid first strand cDNA synthesis kit, were obtained from Thermo Fisher Scientific. TRITC conjugated phalloidin, DAPI, Cultrex™ spheroid invasion matrix were bought from Sigma Aldrich. Oligonucleotides for semi-quantitative RT-PCR were procured from Eurofin, India. GoTaq® Green PCR master mix was obtained from Promega. All other chemicals used were of cell culture grade and procured from Merck and Himedia. All regular plastic wares were of tissue culture grade and brought from Tarsons and Nest.

### 2 Cell culture

MCF7 and MDA MB 231 cell lines were procured from NCCS, Pune, India and maintained in DMEM supplemented with 10% fetal bovine serum and 1% penicillin-streptomycin solution. Human mesenchymal stem cells were a generous gift from Dr. Geetanjali Tomar, IBB. Cells were grown in humidified CO_2_ incubator at 37°C for 2-3 days to reach confluency. Cells were further harvested using Trypsin-EDTA (0.25%) solution and counted before the fabrication of multi-cellular spheroid.

### 3 Nutritional, Hypoxic and Combined stress

Spheroids were formed under normal conditions as stated earlier ^36^. Briefly, 5000 cells/well were seeded on agarose (1%) layered multi well plate and centrifuged at 1200 g for 3 min. After incubation under normal cell culture condition for 3-5 days spheroids of 150-200 µm diameters were formed.

Spheroids were treated with different combinations of glucose and serum [**Control**: 4.5 gm/lt glucose+10% serum; **H-5**: 4.5 gm/lt glucose+5% serum; **H-0**: 4.5 gm/lt glucose+0% serum; **L-5**: 1 gm/lt glucose+5% serum; L-0: 1 gm/lt glucose+0% serum] to create nutritional stress conditions. Cells cultured in 2D condition were treated with CoCl_2_ (200 µM) to create pharmacological hypoxia (**Hy**) and confirmed by expression of hypoxia related marker gene (Hif1). Combined stress conditions are mentioned as H-5-Hy, H-0-Hy, L-5-Hy and L-0-Hy throughout the text.

### 4 Acute and chronic stress conditions

Nutritional and hypoxic stresses were induced individually and in combined conditions to create multiplex stress conditions. Exposure to these individual and multiple stress conditions was further controlled and coined as acute and chronic stress. For acute stress, the fabricated spheroids were harvested and exposed to different stress conditions (individual and combined) for 72 hr. For chronic stress, cells were subjected to spheroid formation protocol directly different stress conditions (individual and combined) for 72 hrs. As the cells were unable to form spheroids under the combined chronic condition, Cultrex™ spheroid invasion matrix was used to fabricate the spheroid ^32^. Stressed spheroids were further collected and used for subsequent experiments.

### 5 Analysis of spheroid growth

Spheroid morphology and proliferation was analyzed using phase contrast microscopy (Nikon Eclipse TiU) after growing them in acute and chronic stress. Aspect ratio of the spheroids was measured using NIS software. The values were plotted using box and whisker plot in Origin Lab software.

### 6 mRNA profiling

Control and stressed MCF7 spheroids were treated with Trizol-chloroform to isolate RNA. Isolated RNA (1 µg) was further used to prepare cDNA following the manufacturer’s protocol (Revertaid first strand cDNA synthesis kit). Equal amount of cDNA was used to assess the mRNA (detail in SI table 1) profile through semi-quantitative RT-PCR using GoTaq® Green master mix and quantified through agarose gel electrophoresis. β-2 microglobulin was used as internal control. Briefly the fold change values were calculated by normalization of each mRNA expression level using internal control through the gel analysis plugin of ImageJ. Normalized values above and below of 1.0 were considered as up and down regulation respectively. The fold change values were plotted using the Heat Map option from the Origin Pro. General colour coding was from Blue to Red depicting the down to up regulation.

Additionally the normalized fold change values of mRNAs of the 23 genes were submitted to Reactome.org database for analysis. All of the entities were found to identify 388 pathways which were hit by at least one of them. To understand the effect on pathway network, genome-wide overview was also generated.

### 7 Analysis of nuclear distribution

Control and stressed spheroids (acute and chronic individual stress) were washed with 1X PBS and fix with 2-4% PFA followed by the permeabilization with 0.01% Triton X100 followed by staining with DAPI for 15 min. Nucleus imaging was done with 60x objective of CLSM (Nikon AR1) and the images were processed with ImageJ software. Peripheral (top 2-4 stacks) nucleus area was measured using the plugins from ImageJ and plotted using box and whisker plot in Origin Lab software. Nucleus morphology of the spheroids from the acute and chronic combined stress conditions cannot be studied due to the fragile nature of the spheroids.

### 8 Distribution of actin filaments

Distribution of cytoskeleton in control and stressed spheroids (acute and chronic individual stress) was analyzed by staining the actin filaments. The control and stressed MCF7 spheroids were washed with 1x PBS and fixed with 2-4% PFA followed by the permeabilization with 0.01% Triton X100. The spheroids were further stained with TRITC conjugated phalloidin (1:100 dilutions) for 2 hr, followed by the imaging in CLSM with 60x oil immersion objective. Cytoskeleton analysis of the spheroids from the acute and chronic combined stress conditions cannot be done due to the fragile nature of the spheroids. Thickness of cortical actin was measured by ImageJ as described elsewhere ^47^. Briefly fluorescence intensity of the 8 bit image was measured and fitted using Gaussian function in Origin Lab. FWHM of the fitted peak was used as a measure of thickness at that cross-section. Thickness values (FWHM) were restricted between 0-1000 nm as mentioned in previous reports. Minimum 10 cells per conditions were taken and 5 ROI measurements per cell were done.

### 9 Pseudo 3D migration and differential inhibition of migration

Spheroids grown in acute (individual and combined) and chronic stress (individual) were subjected to pseudo 3D migration under stress (similar as before) and rescued (4.5 gm/lt glucose and 10% serum) condition. The stressed/rescued spheroids were allowed to migrate on cell culture treated glass cover slips/24 well plates for 20-24 hr. Phase contrast images of migrated cells were acquired using phase contrast microscopy and analyzed.

Profiling of secretory matrix metallo proteases of single cell migration conditions (acute L-0, H-5-Hy, H-0-Hy and L-5-Hy) was done by gelatine based zymogram.

The modality of singly migrating cells was identified by differential inhibitor assay. Briefly Marimastat, ML7, and blebbistatin were added to the rescue media (10 µM) and incubated for 24 hr. Number of migrating cells was counted using phase contrast microscopy.

### 10 Spheroid formations in invasion matrix and sprouting assay

Cultrex™ Spheroid invasion matrix was used to fabricate spheroids under chronic combined stress following the manufacture’s protocol, as the agarose plate protocol was not successful. Briefly 3000 cells were seeded into the invasion matrix and centrifuged for 2 min at 200 g. The plate was incubated in 37°C in humidified CO_2_ (5%) incubator for 72 hr to allow spheroid formation to take place under chronic combined stress conditions. Phase contrast images of formed spheroids were acquired through phase contrast microscope. To analyze the migration of these stressed spheroids, laminin (1µM) was added to the invasion matrix along with the rescue media. Formation of sprout-like structures was monitored and images were acquired after 72-96 hr.

### 11 Effect of stressed spheroids on stem cell migration after rescue

Spheroids are fabricated from pre-stained (Cytotracker green™) cancer cells and exposed to acute combined nutritional stress (H-5-Hy, H-0-Hy, L-5-Hy and L-0-Hy) for 72 hours followed by the rescue with fresh media. For the stem cell-tumor interaction assay rescued spheroids were transferred to agarose coated 96 well plates. Each spheroid was further overlaid with 50 µl collagen solution (1mg/ml) followed by NaOH mediated polymerization of the collagen. After the polymerization, pre-stained (Cytotracker red™) stem cells were seeded (10,000 cell/well) on top of the collagen layer. Directional migration of the stem (red) cell towards the underlying spheroids (green) and recruitment were monitored over next 24 hr and imaging was done using CLSM using the Z scanning mode.

### 12 Statistical analyses

All experiments were repeated more than 3 times with minimum three technical replicates. For normally distributed data set, the significance of difference in mean value of the population was analyzed by parametric tests such as one way ANOVA followed by the post hoc testing such Tuke’s test (Origin Lab). Significance level used in all tests was 0.05 and p<0.05 was considered significant.

## Acknowledgement

T.D thanks UGC for the UGC-FRP scheme, DST SERB and DBT for research funding. S.G thanks DST SERB for the fellowship. A.T thanks CSIR, India for the fellowship. A.N thanks UPE II for the fellowship. The authors thank IBB, SPPU and UPE II for infrastructure, Central research facilities and funding respectively. The authors thank Dr. Geetanjali Tomar for providing the human mesenchymal stem cells and Dr. Ranjana Rai for critical reading of the manuscript and suggestions.

## Author Contributions

T.D. designed the experiments, analyzed the data and wrote the manuscript; S.G, A.T and perform the experiments. All authors have given approval to the final version of the manuscript.

