## Supplemental data for "Nutritional stress alone can control *in vitro* tumor formation and its invasive nature" (from another version of this preprint: version 1, 2020-02-17)

SI Table 1 Gene name list used in the mRNA profiling study

| Sr. No | Gene name |
| --- | --- |
| 1 | PHGDH \| |
| 2 | PKC ζ /PKC2 |
| 3 | mTOR |
| 4 | AMPK |
| 5 | HIF1 alpha/MOP1 |
| 6 | VHL |
| 7 | CA9 |
| 8 | AGTR/AT1 |
| 9 | PDGFRβ |
| 10 | SITR/SIT1 |
| 11 | MMP2 |
| 12 | LOX |
| 13 | FOXA3a/TCF3G |
| 14 | Cathepsin/CTSB |
| 15 | CD144/CDH5 |
| 16 | TCF7 |
| 17 | Axin2 |
| 18 | β-catenin/CTNNB1 |
| 19 | Integrin/ITGB1 |
| 20 | EGFR |
| 21 | ERK2/PRKM1 |
| 22 | P21/CAP20 |
| 23 | PKCD/PRKCD |

SI-Table 2

The table shows the 25 most relevant pathways involving the 23 candidate genes identified by Reactome database sorted by p-value.

| Pathway name | Entities |
| --- | --- |
| Binding of TCF/LEF:CTNNB1 to target gene promoters | 4/10 |
| Interleukin-4 and Interleukin-13 signalling | 8/211 |
| Signalling by receptor Tyrosine Kinases | 11/554 |
| Signalling by Interleukins | 11/639 |
| Signalling Transduction | 23/3305 |
| Cytokine signalling in Immune system | 14/1261 |
| Regulation of gene expression by Hypoxia-inducible Factor | 3/15 |
| Repression of WNT target genes | 3/16 |
| Transcriptional regulation of AP-2 (TFAP2) family of transcription factors | 4/52 |
| Immune system | 20/2822 |
| VEGFA/VEGFR2 Pathway | 5/126 |
| Signalling by VEGF | 5/135 |
| Formation of beta-catenin: TCF transactivating complex | 4/67 |
| Signal transduction by L1 | 3/25 |
| Ca2+ pathway | 4/81 |
| Cellular response to hypoxia | 4/85 |
| Degradation of beta-catenin by destruction complex | 4/91 |
| TFAP2(AP2)family regulates transcription of cell cycle factors | 2/6 |
| Transcriptional activation of p53 responsive gene | 2/6 |
| Transcriptional activation of cell cycle inhibitor p21 | 2/6 |
| Energy dependent regulation of mTOR by LKB1- AMPK | 3/36 |
| RHO GTPases Activates NADPH Oxidases | 3/38 |
| RUNX3 regulates CDKN1A transcription | 2/8 |
| Extra-nuclear estrogen signalling | 4/110 |
| VEGFR2 mediated vascular permeability | 3/44 |

SI Table 3

Tabular depiction of spheroid fabrication condition, migration assay conditions and migration pattern

| *Spheroid grown in* | *Migration condition* | *Migration modality* | | | |
| --- | --- | --- | --- | --- | --- |
|  |  | *No migration* | Collective | Single cell | Mixed |
| *Acute* | Stress | H-0, L-0 | H-5, L-5, Hypoxia | - | - |
|  | Rescued | - | H-5, H-0, L-5, Hypoxia | L0 | - |
| *Chronic* | Stress | H-5, H-0, L-5, L-0 | Hypoxia | - | - |
|  | Rescued | H-0, L-0 | H-5, L-5, Hypoxia | - | - |
| *Acute combined* | Stress | H-0-Hyp, L-0-Hyp | H-5-Hy, L-5-Hyp, | - | - |
|  | Rescued | L-0-Hyp | - | - | H-5-Hyp, H-0-Hyp, L-5-Hyp |

SI Fig 1

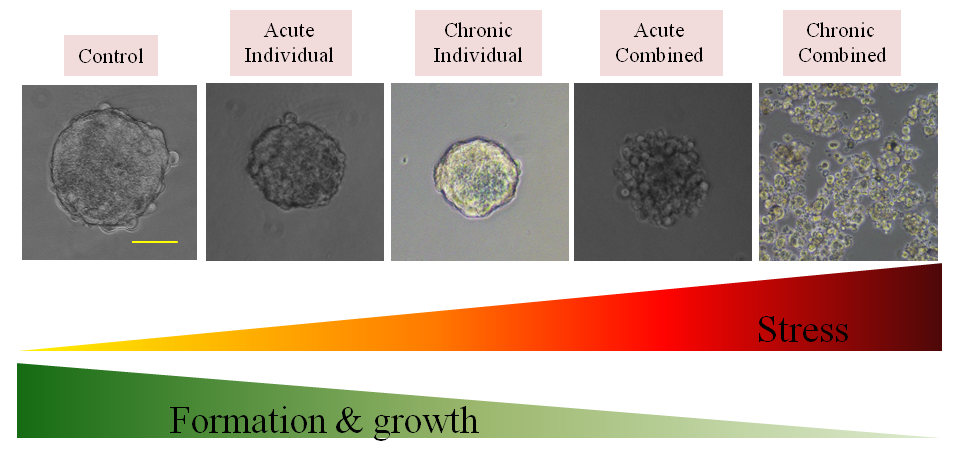

Graphical representation of the different stress levels used in the study and its impact on the spheroid formation, morphology, and size. Scale bar is 10 µm

SI Fig 2

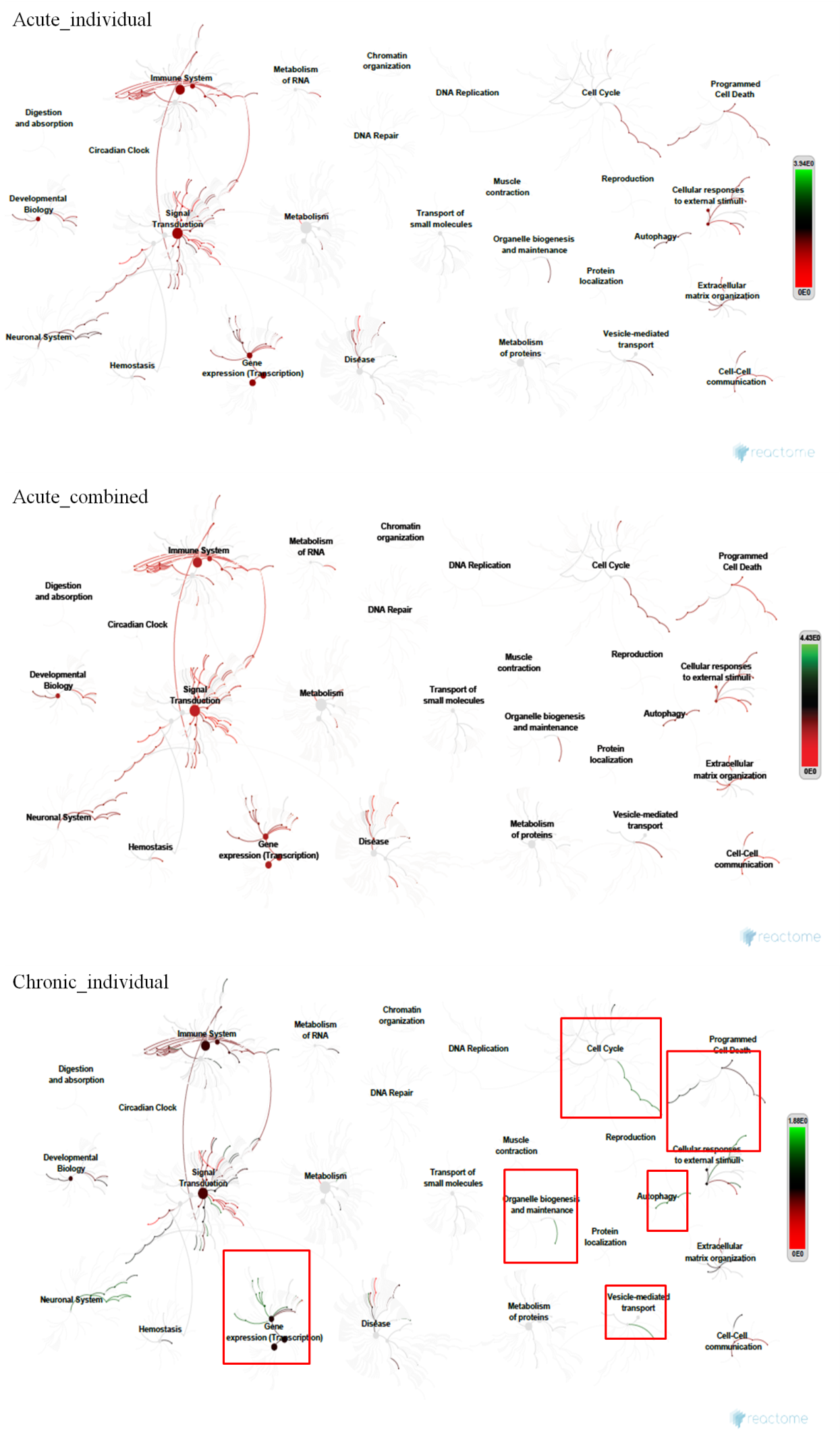

Genome-wide overview of the pathway analysis of 23 genes studied in the report. The color code denotes over-representation of that pathway in the input dataset. Light grey signifies pathways which are not significantly over-represented, where as red pathways represent active pathways, followed by green ones which are over-expressed.

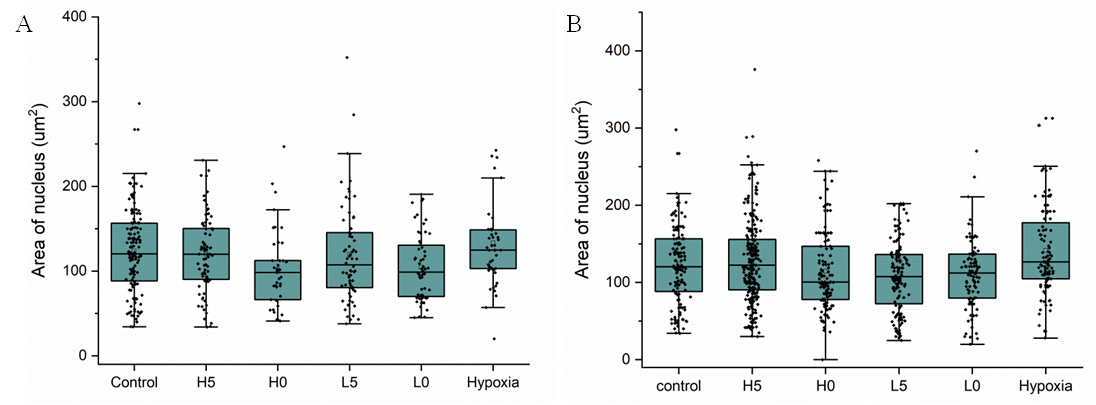
SI Fig 3

Surface area of nucleus measured from top 1-2 cell layers within spheroids stressed acutely (A) and chronically (B). Significance level used in all tests was 0.05 and p<0.05 was considered significant.

SI Fig 4

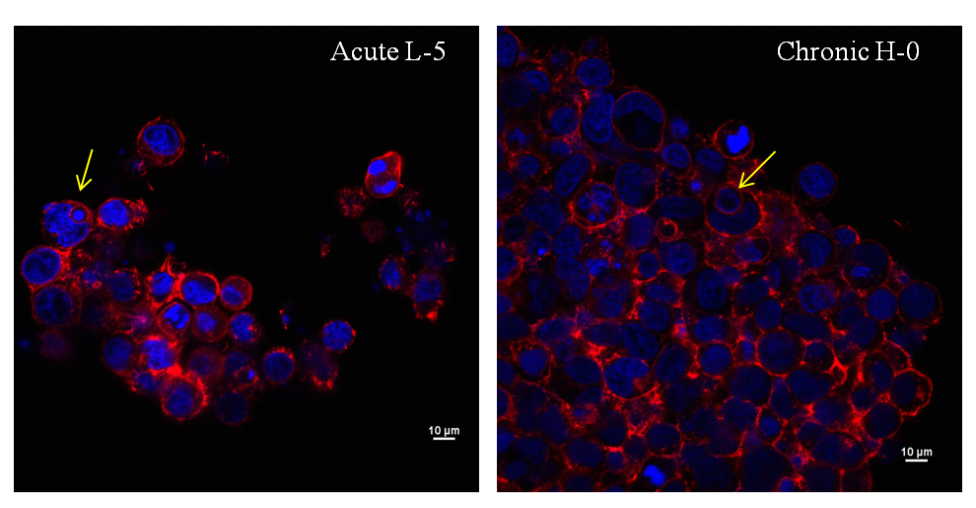

Analysis of cell in cell phenomena in stressed spheroids**.** Spheroids under L-5 (acute) and H-0 (chronic) conditions were subjected to fixation (4% PFA), permeabilization (0.01% TritonX100) and staining with Rodamine conjugated phalloidin and DAPI. Imaging was done in CLSM (60x objective) in Z scanning module. Scale bar is 10 µm.

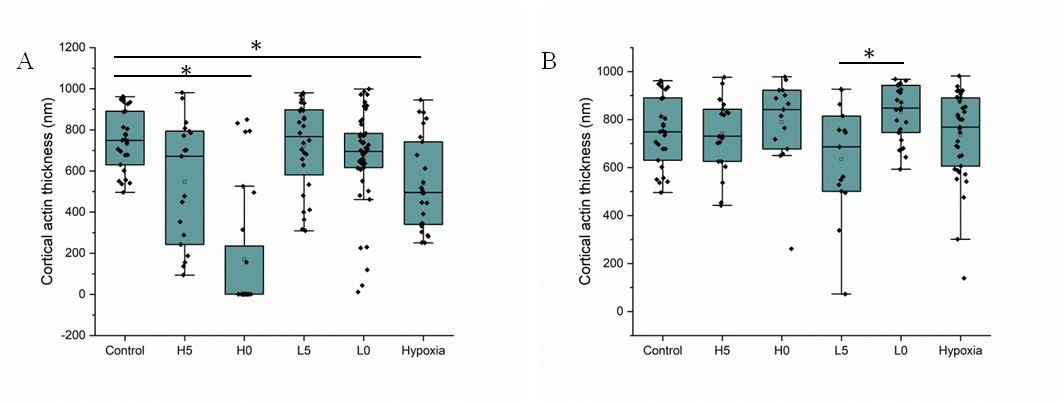
SI-Fig 5

SI-Fig 5: Thickness of cortical actin in control and stressed spheroids stressed acutely (A) and chronically (B) measured through FMHW method. Significance level used in all tests was 0.05 and p<0.05 was considered significant.

SI Fig 6

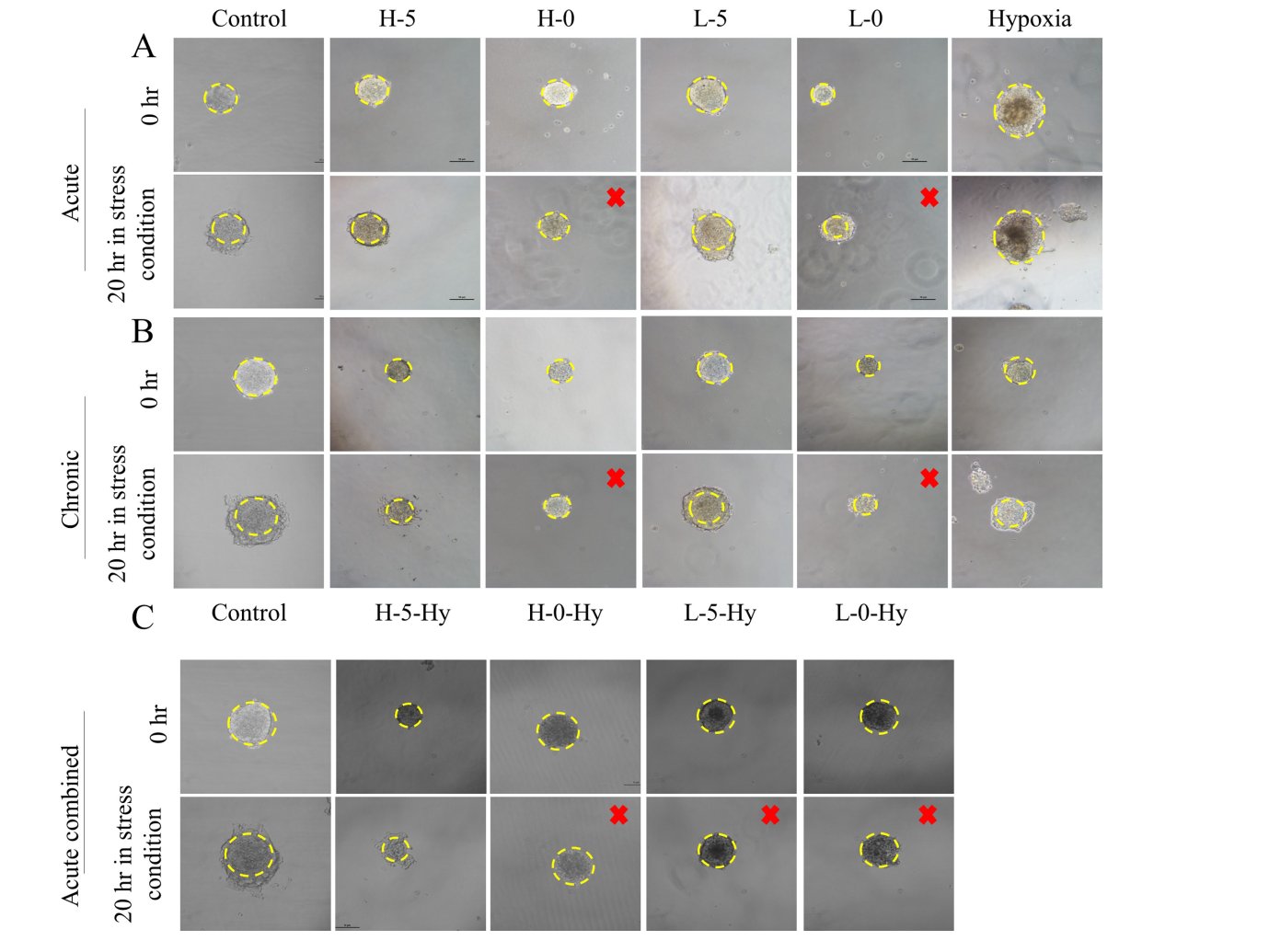

Pseudo 3D migration analysis of the continuous stress (stressed spheroids subjected to respective stress conditions). Most of the spheroids exhibited no migration at all and remained dormant (marked with Red Cross), while rest of them displayed minimal epithelial migration.

SI-Fig 7

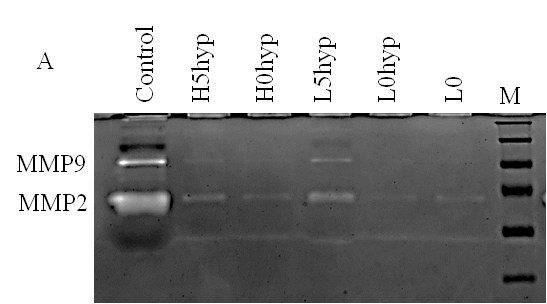

Analysis of secreted MMP (gelatinolytic) by stressed spheroids those exhibit single cell migration after rescue. M denotes marker.

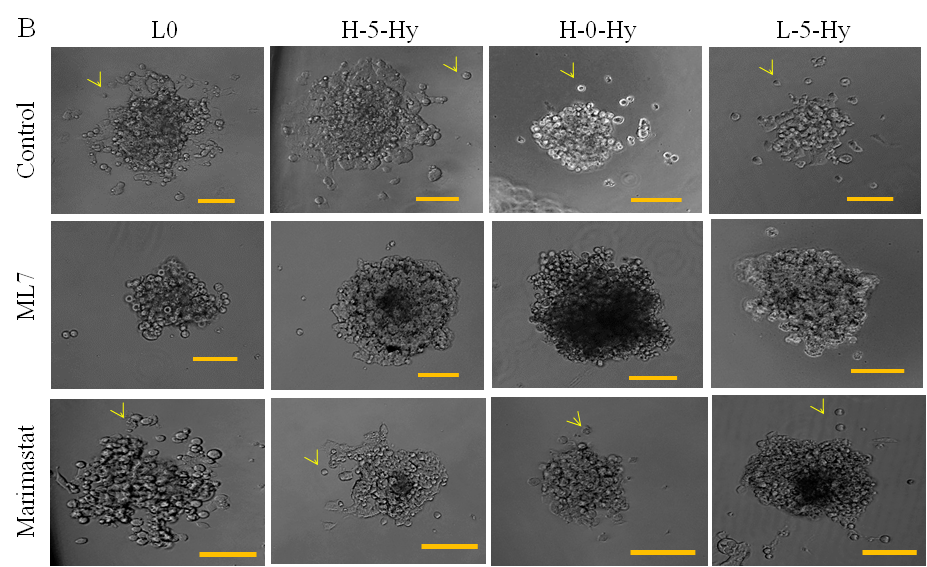

Differential inhibitor assay of pseudo 3Dmigration of severely stressed spheroid under the rescued condition. The singlecells are indicated with yellow arrows in the control panel. After treatment withML7/Marimastat single-cell migration is observed to be completely diminished inML7. Scale bar is 10 µm.
